# Spontaneously Hypertensive Rat substrains show differences in premorbid addiction vulnerability traits and cocaine self-administration: Implications for a novel rat reduced complexity cross

**DOI:** 10.1101/2021.03.06.434216

**Authors:** Kathleen M. Kantak, Carissa Stots, Elon Mathieson, Camron D. Bryant

## Abstract

Forward genetic mapping of F2 crosses between closely related substrains of inbred rodents - referred to as a reduced complexity cross (RCC) - is a relatively new strategy for accelerating the pace of gene discovery for complex traits, such as drug addiction. RCCs to date were generated in mice, but rats are thought to be optimal for addiction genetic studies. Based on past literature, one inbred Spontaneously Hypertensive Rat substrain, SHR/NCrl, is predicted to exhibit a distinct behavioral profile as it relates to cocaine vulnerability traits relative to another substrain, SHR/NHsd. Direct substrain comparisons are a necessary first step before implementing an RCC. We evaluated a number of premorbid addiction vulnerability traits and cocaine self-administration behaviors using a longitudinal within-subjects design. Trait impulsivity and compulsivity were greater in SHR/NCrl than SHR/NHsd, as were reactivity to sucrose reward, sensitivity to acute psychostimulant effects of cocaine, and cocaine abuse liability studied under fixed-ratio and chained schedules of cocaine self-administration. Trait compulsivity correlated with the acute psychostimulant effects of cocaine, which in turn correlated with cocaine taking under the chained schedule. Trait compulsivity also was the best predictor of cocaine seeking responses. Heritability estimates indicated that 22%-40% of the variances for the above phenotypes can be explained by additive genetic factors, providing sufficient genetic variance to conduct genetic mapping in F2 crosses of SHR/NCrl and SHR/NHsd. These results provide compelling support for using an RCC approach in SHR substrains to uncover candidate genes and variants that are of relevance to cocaine use disorders.

**Highlights:**

- Closely related SHR substrains have distinct cocaine vulnerability traits
- Inhibitory control was poorer in SHR/NCrl than SHR/NHsd
- SHR/NCrl were more sucrose reactive and sensitive to acute cocaine than SHR/NHsd
- Cocaine abuse liability was greater in SHR/NCrl than SHR/NHsd
- SHR substrains can be used in an RCC to uncover cocaine vulnerability genes & variants

## 1. Introduction

The addictions, in particular cocaine addiction, are highly heritable neuropsychiatric diseases [1, 2]. Given the nearly 2 million past-month cocaine users in the US, with ∼1 million meeting diagnostic criteria for cocaine dependence [3], new research directed at identifying genetic variants that influence cocaine addiction vulnerability will improve diagnosis, prevention, and treatment. One of the only known genome-wide association study (GWAS) hits to date in cocaine dependence was a variant near *FAM53B* [4] that was functionally supported by covariance between *Fam53b* expression and cocaine self-administration in recombinant inbred mice [5]. To date, human GWAS of SUD, especially those comprising stimulant and opioid use disorders, lack sufficient sample sizes and thus adequate power to detect the small effects of the vast majority of common variants on risk for substance use disorders [6, 7]. Mammalian forward genetic mapping studies can more rapidly achieve appropriate sample sizes and power to identify genome-wide significant genetic loci underlying behavioral traits relevant to the addictions. Nevertheless, a major limitation of many rodent genetic studies is poor mapping resolution, namely those employing F2 crosses, thus leaving investigators with highly significant loci, yet an intractable number of genes within these large intervals (typically 15-20 cM or ∼30- 40 Mb) that contain causal genes and variants. In addition to poor mapping resolution, one must also consider the genetic complexity of loci whereby hundreds of genes and thousands of variants typically underlie F2-derived QTLs between classical inbred strains.

A relatively new approach [8–11] involves forward genetic mapping of complex traits in an F2 cross between closely related substrains of inbred rodents - referred to as a reduced complexity cross (RCC). QTL and fine mapping can be used to resolve quantitative trait loci (QTLs/eQTLs) and quantitative trait genes (QTGs) in genetic crosses that segregate a low level of genetic diversity. Kumar and colleagues used an RCC between C57BL/6 substrains to map and validate Cyfip2 in cocaine sensitivity [12]. We subsequently used an RCC between similar substrains to map the same *Cyfip2* locus and validate *Cyfip2* as a genetic factor underlying binge-like eating [13]. Reduced genetic complexity in C57BL/6 substrains was also used to validate a functional intronic variant in the alpha-2 subunit of the GABA-A receptor in regulating Gabra2 transcript and protein expression via CRISPR/Cas9 gene editing [14]. Finally, Phillips and colleagues exploited reduced genetic complexity between DBA/2 substrains to identify a functional coding variant in Taar1 (trace amine-associated receptor 1) underlying differences in the aversive properties of methamphetamine self-administration and body temperature [15, 16]. These examples show that the use of RCCs for complex trait analysis can accelerate the pace of gene discovery for complex traits relevant to addiction and thus address this challenging public health concern.

The RCC approach to date has only been conducted in mice, but it is widely recognized that rats are optimal for addiction genetic studies because phenotype definitions for addiction vulnerability are well established, readily demonstrated with low attrition rates and more clinically relevant compared to most mouse models [17]. Although several rat models have emerged over the years to evaluate addiction-vulnerable traits such as High vs. Low Novelty Seekers and High vs. Low Impulsive Rats [18, 19], none are as ideal a tool for studying the genetics of addiction vulnerability as inbred Spontaneously Hypertensive Rats (SHR). There is a sizeable literature on drug abuse liability in SHR. We have extensively characterized cocaine self-administration in male SHR/NCrl on several indices of cocaine abuse liability [20–24].

Across studies, we demonstrated that male SHR/NCrl acquired cocaine self-administration faster, exhibited escalated cocaine intake across a range of doses; displayed greater reinforcing strength and motivation for cocaine, and were more reactive to cocaine cues compared to inbred Wistar Kyoto (WKY/NCrl) and outbred Wistar controls. Besides cocaine, male SHR/NCrl self-administered greater amounts of heroin [25] as well as d-amphetamine [26] and methylphenidate [27] compared to WKY/NCrl and/or Wistar. Additionally, male and female SHR/NCrl consumed more ethanol than Sprague-Dawley control rats [28]. These distinct behavioral phenotypes in SHR/NCrl capture several of the salient hallmarks of a substance use disorder outlined in DSM-V [29].

To justify the use of the RCC approach in SHR substrains, it is necessary to first demonstrate a behavioral difference between two closely related substrains. There are at least 8 closely related SHR substrains from different international sources, but none have been directly compared for addiction vulnerability traits. In this report, we phenotyped two substrains – SHR/NCrl (Charles River Laboratories) and SHR/NHsd (Harlan Sprague-Dawley). SHR breeding history began in 1963 at Kyoto University when the outbred Wistar stock in Kyoto (WKY) was selectively bred for elevated blood pressure [30]. Incompletely inbred F13 SHR rats were transferred to the NIH in 1966. SHR rats at NIH were subsequently fully inbred by 1969 and then distributed from NIH (SHR/N) to Charles River Labs (SHR/NCrl) in 1973 at F32 and to Harlan (SHR/NHsd), but with the date of transfer and filial generation at transfer not publicized. Critically, male SHR/NHsd did not self-administer greater amounts of d-amphetamine and methylphenidate [31, 32], and female SHR/NHsd did not self-administer greater amounts of low-dose nicotine than WKY/NHsd, Wistar, and/or Sprague-Dawley controls [33]. The only exception to these distinct substrain profiles is that male and female SHR/NHsd did self-administer greater amount of high-dose nicotine compared to WKY/NHsd controls [33, 34]. In summary, a preponderance of circumstantial evidence predicts that SHR/NCrl are likely to exhibit a distinct behavioral profile as it relates to drugs of abuse compared to SHR/NHsd; however, to our knowledge, these substrains have yet to be compared directly at the behavioral level.

Whole genome sequencing studies demonstrated that SHR/NCrl and SHR/NHsd are closely related (<0.5% genetic variation) and exhibit comparable elevated blood pressure at ∼ 7 weeks of age [35]. SHR/NCrl and SHR/NHsd are viewed as distinct substrains based on analyses from a 10K DNA microarray showing 2.22% heterozygous single nucleotide polymorphisms in SHR/NHsd vs. 0.02% to 0.17% heterozygous single nucleotide polymorphisms in SHR from Charles River Laboratories and the other international sources [36]. Such genetic divergence in SHR/NCrl vs. SHR/NHsd could cause substrain differences in behavior. Thus, in the present study, we directly compared SHR/NCrl and SHR/NHsd substrains on several behavioral indices related to cocaine addiction vulnerability. The phenotypic differences that we report in this study will be exploited to advance our future goal of applying an RCC approach in SHR substrains to identify the genetic basis of variability in cocaine addiction vulnerability traits.

## 2. Materials and methods

### 2.1 Animals

Ten male SHR/NCrl (8 weeks old on arrival; 150-225 g; Charles River Laboratories, USA) and ten male SHR/NHsd (8 weeks old on arrival; 175-250 g; ENVIGO, USA) were housed individually in ventilated cages under a 12hr light/dark cycle (08:00hr on; 20:00hr off) in a climate-controlled vivarium. During food-motivated procedures for trait impulsivity and trait compulsivity, rats were maintained at 80-85% of their expected free-feeding body weight by restricting food in their home cages and providing free access to water. During the remaining procedures, rats had unlimited access to food and water in their home cages. All procedures complied with the 8th edition of the NIH Guide for Care and Use of Laboratory Animals and were approved by the Boston University Institutional Animal Care and Use Committee.

### 2.2 Apparatus

Operant conditioning chambers (model ENV-008CT; Med Associates, St. Albans, VT, USA) were used for tasks that measured trait impulsivity, trait compulsivity and cocaine abuse liability. Each operant chamber was outfitted with two retractable levers, two white stimulus lights, a house light, a speaker, a pellet dispenser, and a syringe pump that were arranged as previously described [37]. Sucrose preference was measured in the rat’s home cage and locomotor activity following acute cocaine administration was measured in boxes containing 15 infrared photobeam detectors (model ENV-3013, Med Associates, St Albans VT, USA). The operant chambers and the locomotor activity boxes were enclosed in sound attenuating cubicles with an exhaust fan.

### 2.3 Procedures

#### 2.3.1 Drugs

Cocaine hydrochloride was obtained from the National Institute on Drug Abuse Drug Supply (Bethesda, MD, USA). For acute intraperitoneal (i.p.) injections (1 ml/kg), cocaine (15 and 20 mg/kg) was dissolved in sterile saline. For chronic intravenous (i.v.) delivery, rats self-administered 0.25 mg/kg cocaine (1.35 mg/ml infused at 1.8 ml/min for 0.6 s/100 g body weight). This is considered a moderate i.v. dose of cocaine, based on its relative position on the descending limb of the fixed-ratio 1 dose-response curve and its relative position on the ascending limb of the progressive-ratio breakpoint dose-response curve in SHR/NCrl, WKY/NCrl and Wistar rats [20, 21, 23].

#### 2.3.2 Catheter Surgery and Maintenance

Prior to initiating drug self-administration sessions, rats were implanted with a catheter made from silicon rubber tubing (Dow Silicones Corporation, Midland, MI, USA; inner diameter, 0.51 mm, outer diameter, 0.94 mm) into the right jugular vein as previously described [37].

Catheters were maintained daily (Monday - Thursday) with 0.1 ml of a saline locking solution containing 100 mg/ml cefazolin (WG Critical Care, LLC, Paramus, NJ, USA) and 30 IU/ml heparin (SAGENT Pharmaceuticals, Schaumburg, IL, USA). Over weekends and holidays, rats received 0.05 ml of a glycerol locking solution containing 3 parts glycerol (Sigma-Aldrich, St.

Louis, MO, USA) to 1 part of the saline locking solution to fill the catheter dead space. Prior to the next behavioral session, the glycerol locking solution was removed and replaced with 0.1 ml of saline containing 3 IU/ml heparin. Catheters were checked daily for leaks and tested periodically for patency by infusing 0.1 ml of a 10 mg/ml solution of methohexital sodium (Brevital; PAR Pharmaceuticals, Chestnut Ridge, NY, USA). Rats received at least 7 days of post-surgical recovery before beginning self-administration sessions.

#### 2.3.3 Trait Impulsivity

Differential reinforcement of low-rate responding (DRL) procedures were implemented to measure trait impulsivity in SHR/NCrl (n=10) and SHR/NHsd (N=10) substrains, as previously described [38]. Briefly, rats first were trained to press the right and left levers under a fixed-ratio (FR) 1 schedule until they reliably obtained 100 food pellets (45 mg chocolate-flavored; Bio-Serv, Frenchtown, NJ, USA) in less than 30 min (typically reached within 2-3 days of training).

Rats were then required during daily (Monday - Friday) 55-min sessions to press a randomly assigned active lever (right or left, counterbalanced across rats) under a DRL 5s schedule with an adjusting limited hold (10s maximum) for a minimum of 10 sessions and until stable responding was reached (<20% change in inter-response-time (IRT) at the 5s wait time for three consecutive sessions). An incremental training protocol was used to progress rats to the DRL 30s schedule that also continued for a minimum of 10 sessions and until stable responding was reached at the 30s wait time for three consecutive sessions. Under DRL 5s and 30s contingencies, responses on the active lever greater than 5s or 30s apart, respectively, were reinforced by 45 mg food pellets, while premature responses reset the 5s or 30s timer. Inactive lever responses were recorded but had no programmed consequences. DRL procedures were conducted in darkened chambers. Measures of impulsive action under DRL responding included response efficiency (percentage of reinforced responses) and burst responding (percentage of active lever responses with IRTs less than 2s apart), which were averaged over the last 3 sessions of stable responding in individual rats under the DRL 5s and DRL 30s schedules.

#### 2.3.4 Trait Compulsivity

Following completion of the DRL procedures, schedule-induced polydipsia (SIP) procedures were implemented to measure trait compulsivity in SHR/NCrl (n=10) and SHR/NHsd (N=10) substrains as previously described [39]. Briefly, the left lever was retracted and the right lever was replaced by a panel accommodating a 100 ml graduated cylinder with 1-ml volumetric markings. The 3-inch ballpoint sipper tube with a 1-inch bend (Ancare Corp., Bellmore, NY, USA) protruded 3.6 cm into the chamber. Before each session, rats were weighed and the graduated cylinders filled with fresh tap water. After attaching the graduated cylinder to clips on the outside of the panel, the initial water level was recorded to the nearest ml. During 60-min daily sessions (Monday – Friday), 45 mg food pellets were delivered noncontingently under a fixed time (FT) 60s schedule. When each session ended, rats were removed immediately from the chamber and the final water level was recorded to the nearest ml and is reported as ml/kg body weight. Rats underwent 12 SIP sessions conducted in chambers illuminated by the house light.

#### 2.3.5 Sensitivity to Sucrose Reward

Following completion of the SIP procedure, rats returned to *ad libitum* feeding for 1 week and then sucrose preference testing was implemented to measure sensitivity to a non-drug reward in SHR/NCrl (n=10) and SHR/NHsd (N=10) substrains as previously described [40].

Briefly, testing was conducted in the rat’s home cage in the animal facility over a 4-day period. On day 1, each rat was provided with two water bottles located on the sides of the central food hopper to habituate rats to drinking from 2 bottles for 23hr. On day 2, one bottle was randomly switched to contain 0.8% sucrose solution midway through the light cycle (12:00hr) to habituate rats to the novel sucrose solution for 23hr. On day 3, the bottles were reversed to avoid perseveration effects and sucrose preference was measured after 23hr. On day 4, the 0.8% sucrose solution bottle was replaced with water, and water preference was measured for 23hr before the two bottles were removed from the cage. The ml/kg consumed from each bottle were recorded on each habituation and test day. Sucrose preference was calculated by dividing the ml/kg sucrose consumed by the total ml/kg sucrose + water consumed on day 3. Water preference was calculated similarly by dividing the ml/kg water consumed from the left bottle by the total ml/kg water consumed from the left and right bottles on day 4. Reactivity to sucrose reward was measured by comparing total fluid intake during the water and sucrose preference tests.

#### 2.3.6 Sensitivity to the Acute Psychostimulant Effects of Cocaine

Following the sucrose preference test, we assessed locomotor activity before and after a cocaine challenge to determine sensitivity to the acute psychostimulant effects of cocaine in SHR/NCrl (n=10) and SHR/NHsd (N=10) substrains. On day 1, rats were placed into the apparatus for a 30-min habituation period and then received an ip injection of sterile 0.9% saline (1 ml/kg) and were returned to the apparatus for 1hr. On days 2 and 3, the procedure was identical except that rats received an ip injection of 15 mg/kg and 20 mg/kg cocaine, respectively. Locomotor activity counts (3-consecutive photobeam breaks) were recorded each day in 5-min bins for the habituation and test phases of each session.

#### 2.3.7 Cocaine Abuse Liability

Catheters were surgically implanted as described in section 2.3.2. Procedures adapted from [41] were used to measure cocaine self-administration behavior in SHR/NCrl (n=6) and SHR/NHsd (n=10) substrains under fixed-ratio and chained schedules.

##### Fixed-Ratio Schedule

During daily sessions (Monday – Friday), rats were randomly assigned a taking lever (counterbalanced left or right across rats) and each press on this lever (FR1) produced a cocaine infusion (0.25 mg/kg) followed by its retraction. The house light then extinguished and the stimulus light above the taking lever was illuminated for the duration of a 20s timeout (TO) period. After the TO, the taking lever was reinserted into the chamber and the house light was re-illuminated. Individual sessions ended after 30 drug infusions or 2hr, whichever occurred first. Rats received a minimum of 10 taking sessions and until responses were stable for three sessions (range 10-11 sessions). The number of taking responses, averaged over the last 3 sessions of stable responding in individual rats, was used to evaluate cocaine taking behavior.

##### Chained Schedule

A chained schedule then was used and each cycle of the seek-take chain started with insertion of the seeking lever, with the taking lever retracted. The first lever press on the seeking lever (FR1) initiated a random interval (RI) schedule and the first lever press made after the RI elapsed retracted the seeking lever and inserted the taking lever. The schedule on the seeking lever was increased from RI 2s to RI 120s over 5 to 10 sessions while the schedule on the taking lever remained at FR1 with a 20s TO after the drug infusion. The post-infusion TO then increased from 20s to 600s over the next 5 to 10 sessions. At the terminal schedule, rats were responding under a chained schedule denoted FR1, RI 120s; FR1, 600s TO. Individual sessions ended after 11 seek-take cycles were completed or 2hr, whichever occurred first. Rats continued with seek-take sessions at the terminal schedule until seeking lever responses were stable for three sessions (range 4-17 sessions at the terminal schedule). The number of seeking responses and number of cycles completed, averaged over the last 3 sessions of stable responding in individual rats, were used to evaluate cocaine seeking and taking under the chained schedule. At the end of behavioral testing, rats were euthanized by an overdose of sodium pentobarbital.

### 2.4 Statistical Analyses

Measures of trait impulsivity (DRL), trait compulsivity (SIP), and sensitivity to sucrose reward (sucrose preference) were analyzed by 2-tailed t-tests for independent samples to compare performances in SHR/NCrl and SHR/NHsd. SIP also was analyzed by 2-factor (substrain X session number) repeated measure analysis of variance (RM ANOVA) to compare the development of polydipsia in SHR/NCrl and SHR/NHsd over the 12 sessions. To evaluate reactivity to sucrose reward in SHR/NCrl and SHR/NHsd, total fluid intake during water and sucrose preference tests was analyzed by 2-factor (substrain X preference test type) RM ANOVA. Locomotor activity in 5-min bins over the 30-min habituation sessions and the 1-hr drug sessions was analyzed by 3-factor (substrain X day/dose X bin) RM ANOVA to measure the time course of basal locomotor activity and the acute psychomotor stimulant effects of cocaine in SHR/NCrl and SHR/NHsd. For drug self-administration data, measures were analyzed by 2-tailed t-tests for independent samples to determine substrain differences in cocaine taking, cocaine seeking, and cocaine seek-take cycles completed. Post-hoc Tukey tests were used following significant ANOVA factors in the above analyses.

The 2-tailed Pearson correlation statistic was used to analyze the associations between selected dependent measures that best characterized the cocaine vulnerability traits of interest: DRL 30s and DRL 5s response efficiency (impulsivity), SIP terminal water consumption (compulsivity), sucrose preference (sensitivity to sucrose reward), total fluid intake during the sucrose preference test (reactivity to sucrose reward), locomotor responses on habituation day 1 (response to novelty), initial 10 min of locomotor responses after 20 mg/kg cocaine (sensitivity to acute cocaine), and cocaine taking, cocaine seeking, and cocaine seek-take cycles completed (cocaine abuse liability). The equation *α*’ = 1 – (1 – overall-*α*)^1/k^ was used to correct for multiple comparisons in the correlation matrix [42]. Based on this equation, probability values <0.031 were considered significant in this study. Narrow-sense heritability (h^2^) was estimated by calculating the total variance, then calculating the average within-strain variance between the two strains (environmental variance), and then subtracting the environmental variance from the total variance which yielded the between-strain variance (genetic). Genetic variance was then divided by the total phenotypic variance which yielded the heritability estimate of the particular trait [43, 44].

## 3. Results

### 3.1 Trait Impulsivity

Under the DRL 5s schedule, response efficiency was similar but burst responding was greater (p<0.05) in SHR/NCrl compared to SHR/NHsd (Figs 1a and 1b). Under the DRL 30s schedule, SHR/NCrl exhibited lower response efficiency (p<0.01) and greater burst responding (p<0.05) than SHR/NHsd (Figs 1c and 1d). The total number of active lever responses under the DRL 30s schedule was greater in SHR/NCrl than SHR/NHsd (375±15 vs. 289±16, respectively; p<0.001), whereas under the DRL 5s schedule the total number of active lever responses did not differ significantly between SHR/NCrl and SHR/NHsd (525±36 vs. 457±30, respectively). Thus, under the DRL 30s schedule for which greater inhibitory control was required, responding was relatively more premature and nonproductive in SHR/NCrl, providing evidence for increased impulsivity.

**Figure 1.**
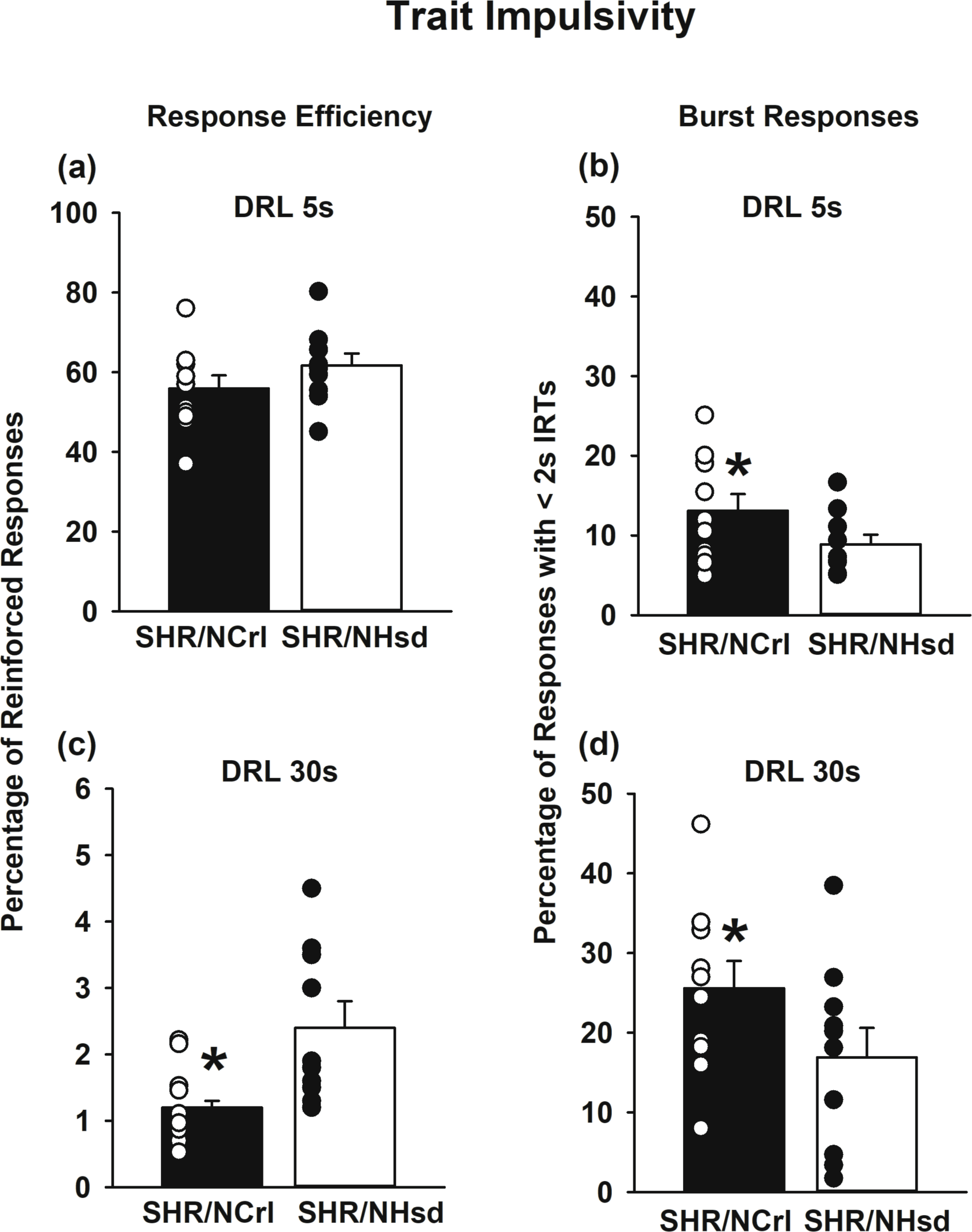
Differential reinforcement of low-rate (DRL) performances in SHR/NCrl (n=10) and SHR/NHsd (n=10) male rats. Values are the mean ± s.e.m. and individual rat data points for the percentage of reinforced responses (response efficacy) and percentage of responses with IRTs < 2s (burst responding) averaged over the last 3 daily sessions at criteria for the DRL 5s wait time (panels a and b) and the DRL 30s wait time (panels c and d). *ps<0.05 comparing SHR/NCrl to SHR/NHsd.

### 3.2 Trait Compulsivity

SHR substrains consumed similar amounts of water at the beginning of the SIP task, but SHR/NCrl developed polydipsia faster and to a greater extent than SHR/NHsd. Across the 12 sessions (Fig 2a), the substrain X session number interaction was significant (F[11, 198]=2.5, p<0.01). Relative to session 1, water consumption increased beginning on session 4 in SHR/NCrl (ps<0.001), but not until session 7 in SHR/NHsd (ps<0.02). Relative to SHR/NHsd, water consumption was greater in SHR/NCrl on sessions 4-8 and 10-12 (ps<0.05). For the terminal three sessions (Fig 2b), there were robust substrain differences between SHR/NCrl and SHR/NHsd (p<0.01). Thus, consummatory behavior in the SIP task was relatively more habitual and excessive in SHR/NCrl, providing evidence for increased compulsivity.

**Figure 2.**
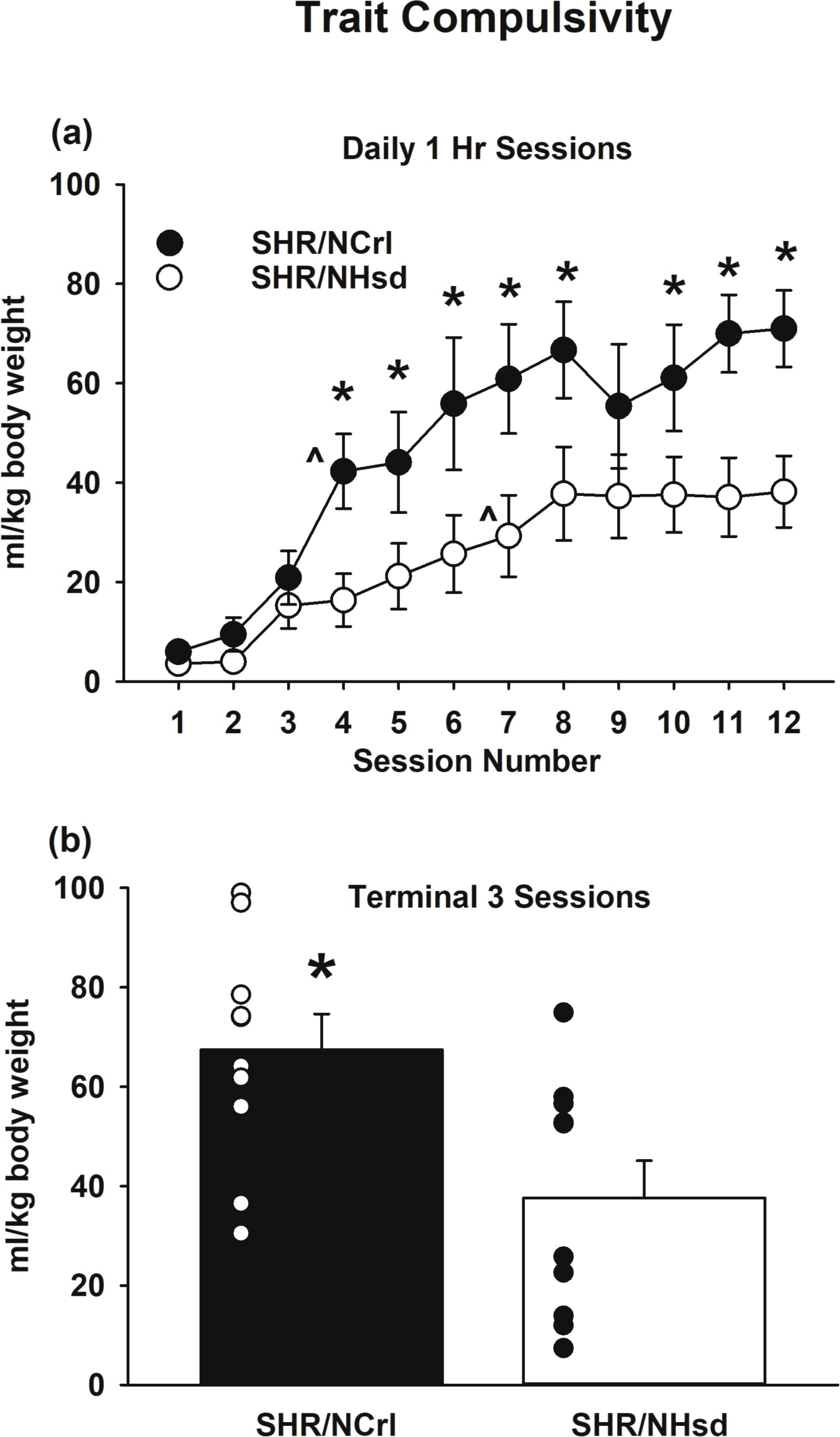
Schedule-induced polydipsia (SIP) in SHR/NCrl (n=10) and SHR/NHsd (n=10) male rats. Values are the mean ± s.e.m. ml/kg water intake for each of the 12 daily sessions (panel a) or the mean ± s.e.m. and individual rat data points for ml/kg water intake averaged over the terminal 3 daily sessions (panel b). * ps<0.05 comparing ml/kg water intake between SHR/NCrl and SHR/NHsd on sessions 4-8, 10-12, and the terminal 3 sessions combined. ^ ps<0.02 relative to session 1.

### 3.3 Sensitivity to Sucrose Reward

The preference ratios for water and sucrose were not significantly different between SHR/NCrl and SHR/NHsd (Fig 3a). Each substrain consumed approximately 50% of their daily fluid intake from each water bottle during the water preference test and approximately 80% of their daily fluid intake from the sucrose bottle during the sucrose preference test. Thus, sensitivity to sucrose reward did not differ between substrains. In contrast, total fluid intake depended on substrain and preference test type (F[1, 18]=5.8, p<0.03). Whereas there were no significant substrain differences in total fluid intake during the water preference test, SHR/NCrl had greater total fluid intake than SHR/NHsd during the sucrose preference test (p<0.001; Fig 3b). Moreover, total fluid intake in SHR/NCrl was greater during the sucrose preference than the water preference test (p<0.004), whereas total fluid intake in SHR/NHsd was statistically similar during the water and sucrose preference tests. Thus, SHR/NCrl were relatively more reactive to sucrose reward.

**Figure 3.**
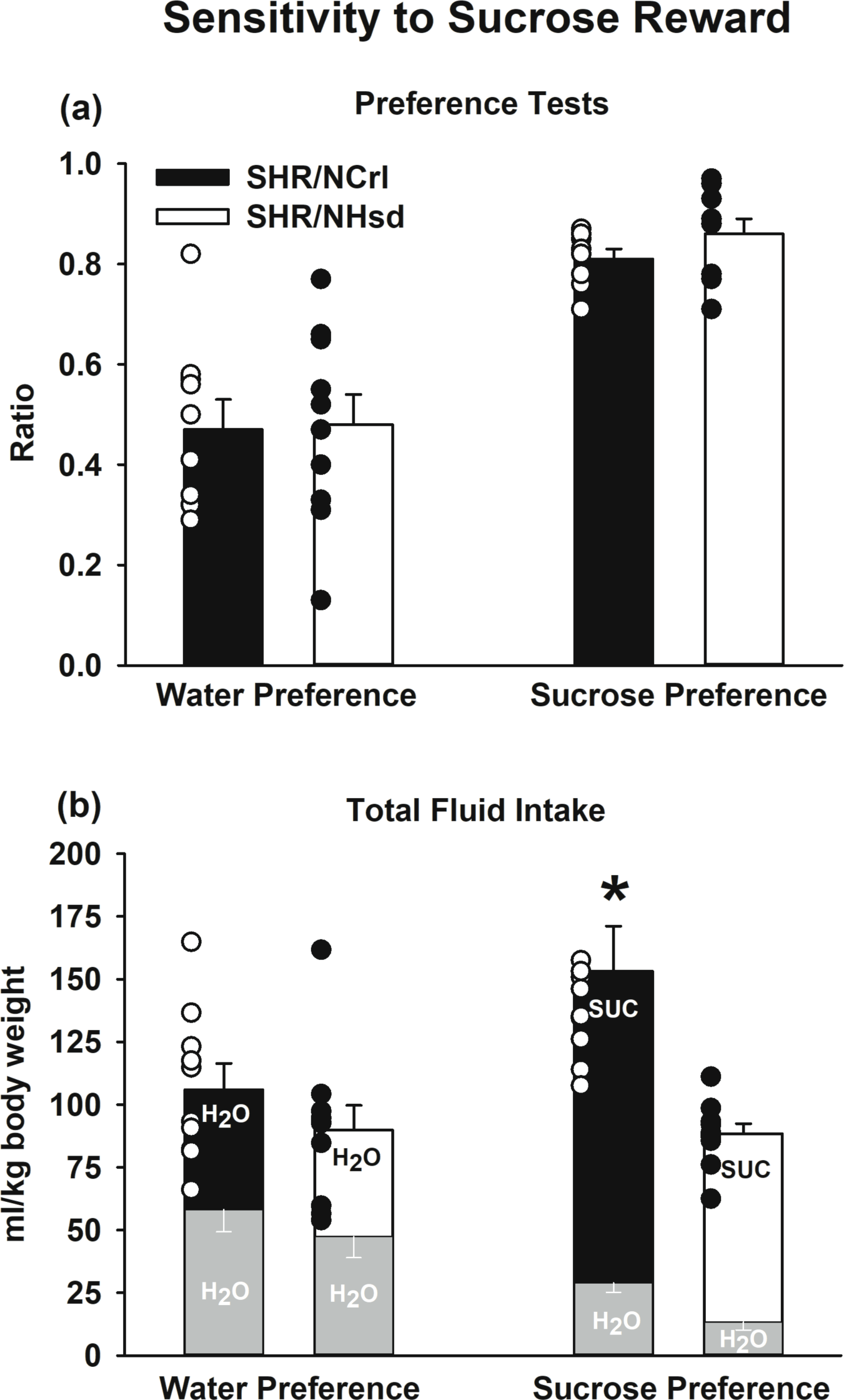
Sucrose preference testing in SHR/NCrl (n=10) and SHR/NHsd (n=10) male rats. Values are the mean ± s.e.m. and individual rat data points for the water and sucrose preference ratios (panel a) and the total ml/kg fluid intake during the water and sucrose preference tests (panel b). * ps<0.004) comparing total intake in SHR/NCrl to SHR/NHsd during the sucrose preference test and comparing total intake during the sucrose preference test to the water preference test in SHR/NCrl.

### 3.4 Sensitivity to the Acute Psychostimulant Effects of Cocaine

During habituation sessions, locomotor activity did not differ significantly between the substrains, but analyses revealed a significant habituation day X bin interaction (F[10, 180]=7.5, p<0.001). Within SHR/NCrl (Fig 4a, left), locomotor activity was greater on habituation day 1 than days 2 and 3 for bins 2-5 (ps<0.03). Within SHR/NHsd (Fig 4b, left), locomotor activity was greater on habituation day 1 than days 2 and 3 for bins 2-4 and 6 (ps<0.01), and on habituation day 1 than day 2 for bin 5 (p<0.001). Thus, there was an overall heightened reaction to the novel environment on habituation day 1, with the response to novelty similar in SHR/NCrl and SHR/NHsd. Basal locomotor activity levels were similar as well between substrains over habituation sessions 1-3.

**Figure 4.**
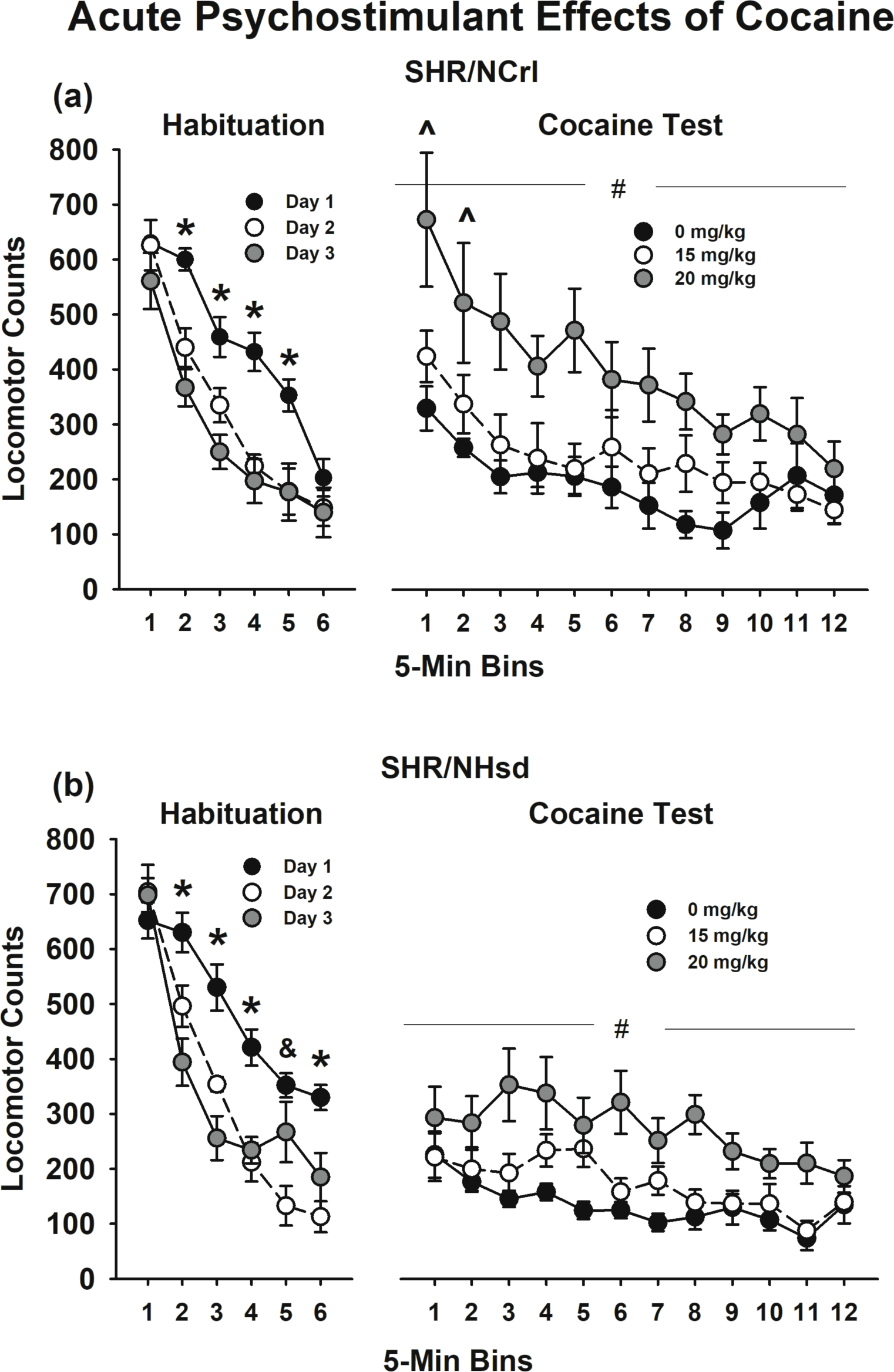
Psychostimulant effects of acute cocaine in SHR/NCrl (n=10) and SHR/NHsd (n=10) male rats. Values are the mean ± s.e.m. locomotor activity counts in 5 min bins over the 30- min habituation sessions and the 1-hr cocaine tests in SHR/NCrl (panel a) and SHR/NHsd (panel b). * ps<0.03 comparing habituation day 1 to days 2 and 3 for bins 2-5 in SHR/NCrl and for bins 2-4 and 6 in SHR/NHsd. & p<0.001 comparing habituation day 1 to day 2 for bin 5 in SHR/NHsd. # p<0.01 comparing 20 mg/kg to 0 mg/kg and 15 mg/kg cocaine in both substrains. ^ ps<0.04 comparing bins 1 and 2 in SHR/NCrl to SHR/NHsd after 20 mg/kg cocaine.

Following acute injections (Fig 4a and 4b, right), cocaine produced dose-dependent increases in locomotor activity (F[2, 36]=19.2, p<0.001). There was greater locomotor activity after 20 mg/kg compared to 0 and 15 mg/kg cocaine within SHR/NCrl and SHR/NHsd (ps<0.01). Analysis of the bin X substrain interaction (F[11, 198]=3.7, p<0.001) revealed that locomotor activity was greater in SHR/NCrl than SHR/NHsd for the first two 5-min bins of the session after 20 mg/kg cocaine (ps<0.04) but not after 0 or 15 mg/kg cocaine. Thus, SHR/NCrl were relatively more sensitive to the acute psychostimulant effects of 20 mg/kg cocaine.

### 3.5 Cocaine Abuse Liability

During initial self-administration training under the FR1 taking schedule (Fig 5a) SHR/NCrl self-administered more cocaine than SHR/NHsd (p<0.05). It should be noted that FR1 training in SHR/NHsd was interrupted between late March and early June 2020 due to a local quarantine related to the COVID-19 pandemic. Rats had just competed the FR1 training phase when the lab was shuttered. At that time, the catheters were filled with the glycerol locking solution, and after its removal in early June, we found that only 1 rat had a nonfunctional catheter that required repair before resuming sessions to repeat the entire FR1 training phase. There were no statistical differences between the initial FR1 baseline (19.5±2.7) and the new FR1 baseline (18.3±3.2) in SHR/NHsd (p<0.49). Thus, the unavoidable interruption in cocaine self-administration during FR1 training did not influence lever responding in SHR/NHsd over the long-term. After transition to the more demanding FR1, RI 120s; FR1, 600s TO chained schedule, SHR/NCrl emitted more cocaine seeking responses (p<0.03) and completed more of the 11 seek-take cycles (p<0.03) compared to SHR/NHsd (Figs 5b and 5c). Thus, SHR/NCrl exhibited relatively greater cocaine abuse liability.

**Figure 5.**
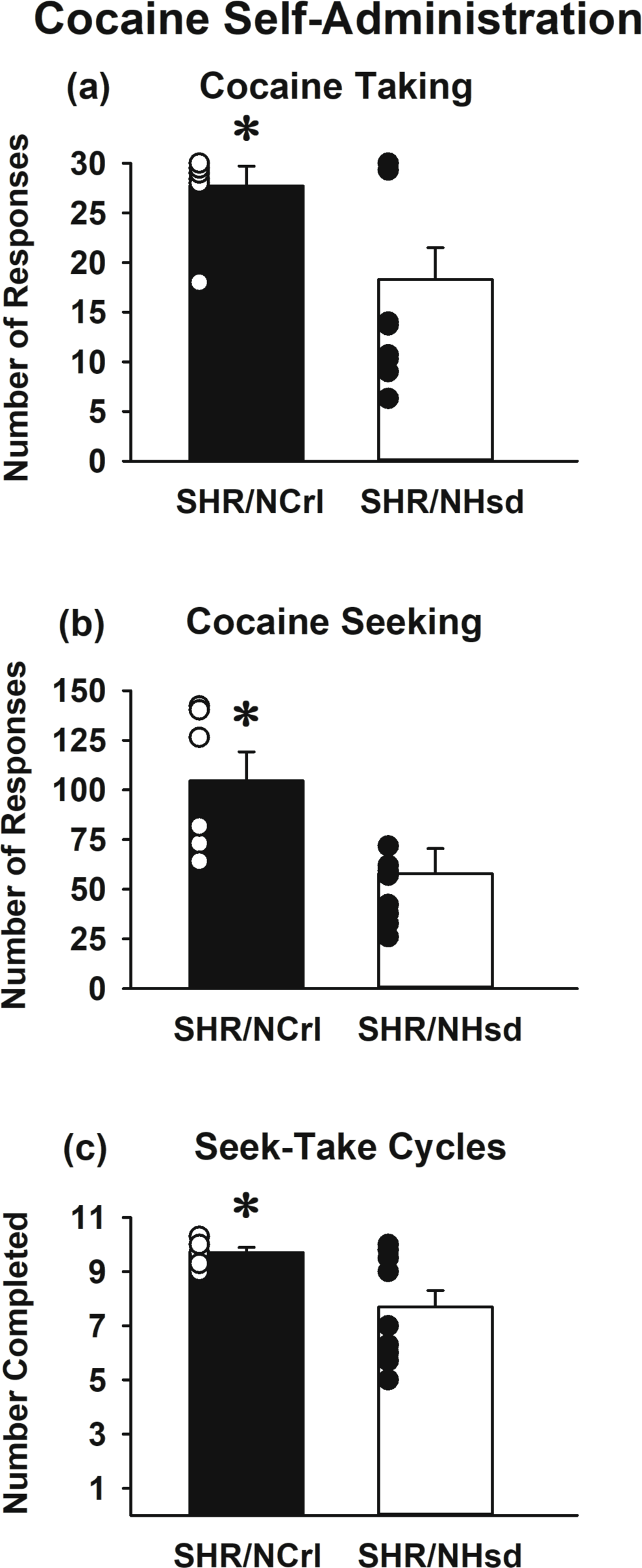
Cocaine self-administration in SHR/NCrl (n=6) and SHR/NHsd (n=10) male rats. Values are the mean ± s.e.m. and individual rat data points for the number of taking lever responses under the FR1 schedule (panel a) and for the number of seeking lever responses (panel b) and seek-take cycles completed (panel c) under the chained FR1, RI 120s; FR1, 600s TO schedule of cocaine (0.25 mg/kg/infusion) delivery. * ps<0.05 comparing SHR/NCrl to SHR/NHsd.

### 3.6 Predictors of Cocaine Abuse Liability and Heritability Analyses

Interestingly, the premorbid level of compulsive behavior exhibited by SHR/NCrl and SHR/NHsd was the best predictor of the acute psychostimulant effects of 20 mg/kg cocaine (Fig 6a) and of cocaine seeking during chronic cocaine self-administration (Fig 6b). Moreover, locomotor activity after acute injection of 20 mg/kg cocaine predicted the number of cocaine seek-take cycles completed (Fig 6c). Cocaine taking behavior under both self-administration schedules predicted the magnitude of cocaine seeking (Fig 6d and 6e). Lower DRL 30s response efficiency (greater impulsivity) was associated with greater compulsive behavior (Fig 6f), while lower DRL 5s response efficiency (greater impulsivity) was associated with greater sucrose reactivity (Fig 6g). Sucrose preference and locomotor response to novelty did not significantly correlate with any behavioral measure evaluated (Table 1).

**Figure 6.**
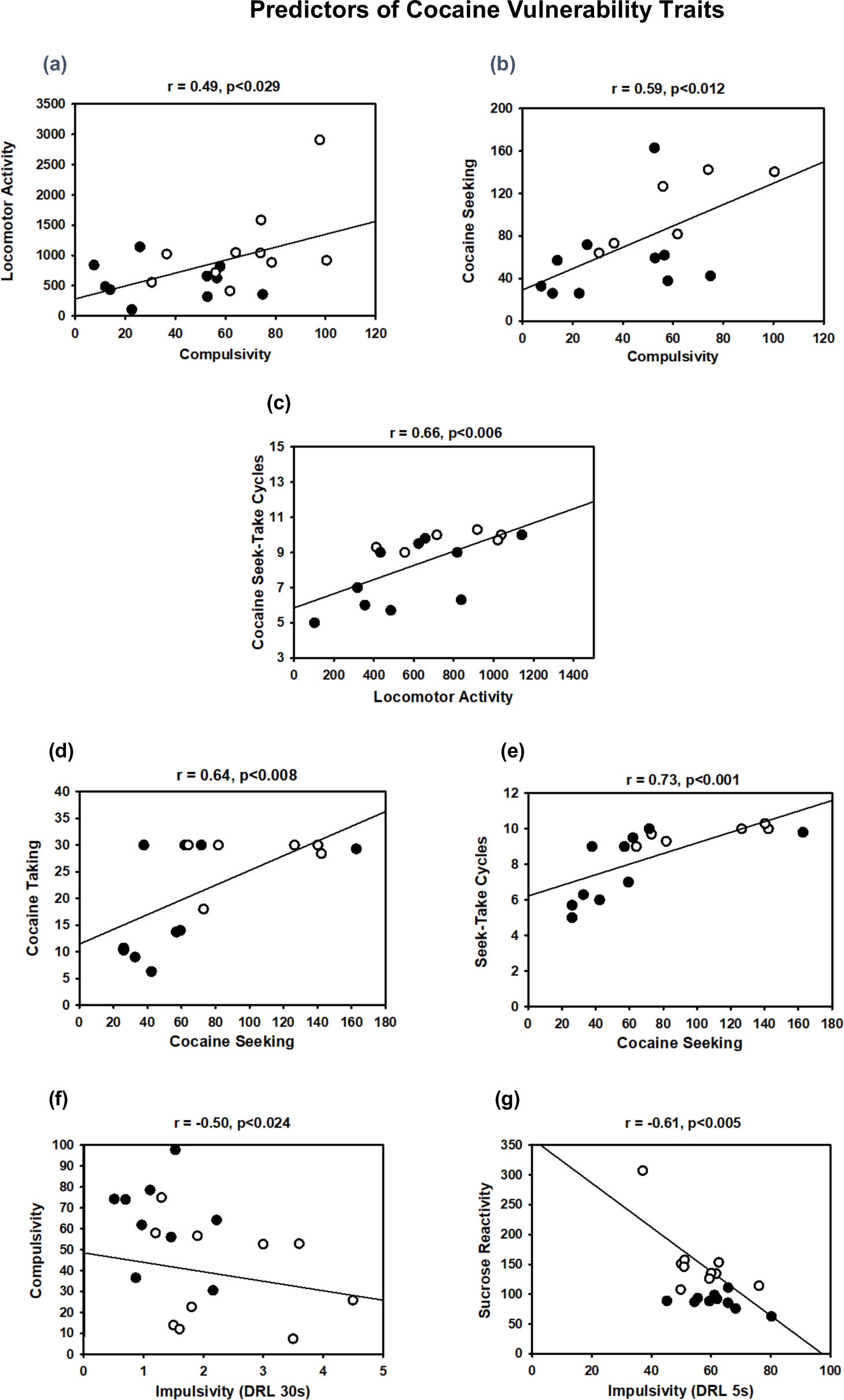
Predictors of cocaine vulnerability traits in SHR/NCrl (white circles) and SHR/NHsd (black circles). Illustrated are behaviors that significantly correlated with trait compulsivity (panels a and b), with cocaine-induced locomotor activity (panel c), with cocaine seeking responses (panels d and e), and with trait impulsivity (panels f and g).

**Table 1.** Correlation matrix for dependent measures in SHR/NCrl and SHR/NHsd substrains. Vertical cell contents: correlation coefficient, 2-tailed p value, and number of observations. Significant correlations are highlighted in gray, after correction for multiple comparisons.

|  | Response<br>Efficiency<br>DRL 30s | SIP | Sucrose<br>Preference | Sucrose<br>Reactivity | Novelty<br>Response | Cocaine<br>Locomotion | Cocaine<br>Taking | Cocaine<br>Seeking | Cocaine<br>Cycles |
| --- | --- | --- | --- | --- | --- | --- | --- | --- | --- |
| Response<br>Efficiency<br>DRL 5s | -0.079<br>0.741<br>20 | -0.062<br>0.794<br>20 | 0.157<br>0.508<br>20 | -0.607<br>0.005<br>20 | -0.083<br>0.729<br>20 | 0.034<br>0.886<br>20 | -0.112<br>0.680<br>16 | -0.294<br>0.269<br>16 | -0.165<br>0.542<br>16 |
| Response<br>Efficiency<br>DRL 30s |  | -0.502<br>0.024<br>20 | 0.053<br>0.826<br>20 | -0.195<br>0.409<br>20 | 0.380<br>0.099<br>20 | -0.136<br>0.568<br>20 | -0.101<br>0.710<br>16 | -0.176<br>0.515<br>16 | -0.169<br>0.532<br>16 |
| SIP |  |  | -0.080<br>0.739<br>20 | 0.143<br>0.548<br>20 | -0.227<br>0.336<br>20 | 0.489<br>0.029<br>20 | 0.455<br>0.077<br>16 | 0.591<br>0.016<br>16 | 0.430<br>0.097<br>16 |
| Sucrose<br>Preference |  |  |  | -0.319<br>0.170<br>20 | 0.046<br>0.846<br>20 | -0.170<br>0.474<br>20 | 0.138<br>0.611<br>16 | -0.130<br>0.632<br>16 | 0.116<br>0.670<br>16 |
| Sucrose<br>Reactivity |  |  |  |  | -0.060<br>0.803<br>20 | 0.225<br>0.340<br>20 | 0.261<br>0.328<br>16 | 0.161<br>0.552<br>16 | 0.200<br>0.457<br>16 |
| Novelty<br>Response |  |  |  |  |  | -0.252<br>0.283<br>20 | -0.119<br>0.661<br>16 | 0.178<br>0.510<br>16 | -0.101<br>0.711<br>16 |
| Cocaine<br>Locomotion |  |  |  |  |  |  | 0.497<br>0.050<br>16 | 0.433<br>0.094<br>16 | 0.659<br>0.006<br>16 |
| Cocaine<br>Taking |  |  |  |  |  |  |  | 0.636<br>0.008<br>16 | 0.862<br>0.00002<br>16 |
| Cocaine<br>Seeking |  |  |  |  |  |  |  |  | 0.728<br>0.001<br>16 |

Narrow-sense heritability estimates (h^2^) were used to determine the percentage of total phenotypic variance explained by additive genetic factors. After estimating heritability based on parental SHR substrain variance, the most heritable phenotype was the number of cocaine seek-take cycles completed (h^2^ = 40%). Other phenotypes showing similar h^2^ estimates were cocaine taking responses (h^2^ = 31%) and reactivity to sucrose reward (h^2^ = 38%). The heritability estimates for DRL 30s impulsivity (h^2^ = 27%) and SIP compulsivity (h^2^ = 27%) were lower and were comparable to the heritability estimates for cocaine seeking responses (h^2^ = 26%). The lowest heritability estimates were for the acute psychostimulant effects of 20 mg/kg cocaine (h^2^ = 16% at the 10-min time point and h^2^ = 22% at the 5-min time point), locomotor response to novelty (h^2^ = 7%), sucrose preference (h^2^ = 5%), and DRL 5s impulsivity (h^2^ = 4%).

## 4. Discussion

### 4.1 Premorbid Inhibitory Control Capacity

Lever responding during the DRL task was more premature and nonproductive in SHR/NCrl than SHR/NHsd, especially for the longer 30s wait time. Lower response efficiency and higher burst responding indicate greater impulsive action [38]. Increased trait impulsivity in SHR/NCrl compared to SHR/NHsd is not surprising, given the historical differences in between- substrain comparisons of behavior during DRL and other tasks of impulsive action or choice.

Collectively, studies showed that SHR/NCrl were consistently more impulsive than both inbred (WKY/NCrl) and outbred (Wistar and Long-Evans but not Sprague-Dawley) control strains [38, 39, 45–50]. In contrast, SHR/NHsd were more impulsive on these tasks compared to one or the other type of control strain, but never compared to both inbred (WKY/NHsd) and outbred (Wistar and Sprague-Dawley) control strains [51–54].

SHR/NCrl were more compulsive than SHR/NHsd in the SIP task in that water intake was more habitual and excessive in SHR/NCrl. By the end of testing, SHR/NCrl consumed ∼65 ml/kg of water and SHR/NHsd consumed ∼40 ml/kg of water in 1hr. Considering that average 24hr water intake in each substrain is ∼80 ml/kg [55, 56], the excessive drinking that developed in 1hr was clearly maladaptive and exceeded the physiological need for water even after consuming 2.7 grams of food during the 1hr test sessions. Individual differences in the development of SIP are considered a means to identify a compulsive endophenotype reflecting poor inhibitory control [57]. To date, SIP has been investigated only in SHR/NCrl for between-strain comparisons, and results consistently showed that polydipsia associated with a FT 60s schedule of food pellet delivery was greater in SHR/NCrl than WKY/NCrl and Wistar control strains [39, 48].

The results of DRL and SIP testing indicate that premorbid inhibitory control capacity is more limited in SHR/NCrl than SHR/NHsd. We chose *a priori* to use the DRL and SIP tasks to study these different facets of inhibitory control because these tasks are conducted under non-overlapping test conditions and have unique requirements, including chamber illumination (dark vs lit, respectively), lever availability (inserted vs. retracted, respectively), basis for food pellet delivery (contingent vs. automatic, respectively), and the behavioral variable assessed (reinforced lever responses vs. ml/kg water intake, respectively). The significant correlation between trait impulsivity and trait compulsivity in this study likely reflects the relationship between the two inhibitory control endophenotypes rather a direct influence of low levels of reinforced lever responses in the DRL task on excessive fluid consumption in the SIP task.

Supporting this assertion was the lack of a significant correlation between low levels of reinforced lever responses in the DRL task and excessive fluid consumption during the sucrose preference test. Heritability estimates indicate that 27% of the variances for trait impulsivity and trait compulsivity can be explained by additive genetic factors. Similar results were reported in a recent twin study showing that inhibitory control traits were heritable, with 33% of the variance for impulsivity and 25% of the variance for compulsivity explained by additive genetic factors [58]. Of relevance, high levels of impulsivity and compulsivity were found to be risk factors for cocaine dependence in both human laboratory studies [59–61] and rat experiments [18, 62, 63]. In the current study, trait compulsivity was a significant predictor of cocaine seeking responses and initial sensitivity to the acute psychostimulant effects of 20 mg/kg cocaine.

### 4.2 Premorbid Reactions to Sucrose, Novelty and Acute Cocaine

Preference for sweet solutions is a measure of sensitivity to a non-drug reward [64].

Sucrose preference did not differ between SHR substrains and heritability was low (5%). Sucrose preference also did not significantly correlate with any cocaine use phenotype that we evaluated, consistent with past research showing that sucrose preference was not correlated with cocaine intake [65]. In contrast, sucrose + water intake during sucrose preference testing (reactivity to sucrose reward) was greater in SHR/NCrl than SHR/NHsd. Past research showed that Sprague-Dawley rats selectively bred for high saccharin intake acquired cocaine self-administration faster and maintained cocaine intake at higher levels compared to Sprague-Dawley rats selectively bred for low saccharin intake [66, 67]. It has been suggested that relatively greater reactivity to sweet solutions in selectively bred rats is a strong indicator of a genetic predisposition for vulnerability to cocaine abuse [68]. The heritability estimate for the sucrose reactivity phenotype indicates that 38% of the variance can be explained by additive genetic factors. This facet of behavior in the SHR substrains might be translational because cocaine-dependent individuals show higher sucrose liking scores than controls [69].

The locomotor response to a novel environment (habituation day 1) did not differ between the SHR substrains, and in support, the estimated heritability was low (7%). Nor did this phenotype significantly correlate with any cocaine use phenotype that we evaluated. This finding is consistent with past research showing that the locomotor response to a novel environment in high and low locomotor responding Sprague-Dawley rats did not predict cocaine intake [19, 70]. Likewise, basal locomotor activity (habituation days 1-3) was comparable in SHR/NCrl and SHR/NHsd. This finding also is consistent with between-strain comparisons showing that SHR/NCrl and SHR/NHsd are similarly hyperactive relative to WKY/NCrl and WKY/NHsd, respectively [38, 51]. In contrast, novelty preference can predict severity of compulsive cocaine use in high and low novelty preferring Sprague-Dawley rats [19], suggesting that novelty preference might be an important phenotype to study in future investigations exploring SHR substrain differences. Previous between-strain comparisons support such efforts, as SHR/NCrl exhibited greater novelty preference than WKY/NCrl and Wistar [26, 71], but novelty preference did not differ significantly between SHR/NHsd, WKY/NHsd and Wistar [31].

The locomotor response to acute 20 mg/kg cocaine was greater in SHR/NCrl than SHR/NHsd for the first 10-min after the injection. This phenotype significantly predicted the number of cocaine seek-take cycles completed under the chained schedule of cocaine self-administration. In a recent study that compared eight inbred mouse strains, the locomotor response to acute 20 mg/kg cocaine depended on strain, with the magnitude of cocaine-induced locomotor activity predictive of cocaine intake and motivation for cocaine self-administration [72]. A study in recombinant inbred mice indicated that 28% to 37% of the variance for locomotor sensitivity to acute cocaine (10-40 mg/kg) can be explained by additive genetic factors [73]. In SHR rat substrains, the heritability estimate was lower and at best, ranged from 22% at 5 min post-cocaine (time point when SHR substrain differences were the greatest) and 16% at 10 min post-cocaine. Notably, initial response to cocaine can be predictive of future cocaine dependence in people [74], supporting our current observation that although not as heritable, cocaine locomotor activity has some predictive value for future cocaine intake. We previously mapped and validated *Hnrnph1* as a quantitative trait gene for methamphetamine-induced locomotor activity and subsequently extended these findings to methamphetamine reward and reinforcement [75–77], providing evidence for at least some portion of shared genetic basis between stimulant-induced locomotor activity and stimulant-induced appetitive behaviors.

### 4.3 Cocaine Abuse Liability

SHR/NCrl self-administered more cocaine than SHR/NHsd under the FR1 schedule and also emitted more cocaine seeking responses and completed more cocaine seek-take cycles under the chained schedule. Heritability estimates indicated that 26% - 40% of the variances for our cocaine use traits can be explained by additive genetic factors, a key finding because cocaine use disorders in people are heritable on the order of 40% - 70% [78, 79]. As discussed in section 4.2, four of our premorbid cocaine vulnerability traits (impulsivity, compulsivity, reactivity to sucrose reward and initial sensitivity to acute 20 mg/kg cocaine) showed substrain differences, with 22% - 38% of their phenotypic variances explained by additive genetic factors. Heritability’s of 20% or greater provide sufficient genetic variance to conduct genetic mapping in experimental crosses [80].

Of the premorbid traits we evaluated in SHR/NCrl and SHR/NHsd, compulsivity and initial sensitivity to acute 20 mg/kg cocaine were the best predictors of cocaine seeking and cocaine taking behaviors, respectively. It was somewhat surprising that impulsivity was not a significant predictor of any aspect of cocaine self-administration behavior in this study, given its prominent association with cocaine self-administration in outbred Lister Hooded rats [18, 62, 81, 82] and outbred Wistar rats [83, 84]. Although both inhibitory control traits were heritable and were correlated in SHR/NCrl and SHR/NHsd, the compulsivity phenotype might be relatively more important for predicting a cocaine-vulnerable phenotype in an F2 population of rats that segregate SHR/NCrl and SHR/NHsd alleles. Interestingly, the balance between brain circuits that promote (ventral striatum-orbitofrontal cortex) and limit (ventral striatum-anterior cingulate cortex) compulsive behavior is dysregulated in cocaine users, with the degree of circuit dysregulation positively correlated with the number of DSM-IV compulsive drug use symptoms [85]. Strikingly similar brain-behavior relationships were reported in a subset of outbred Sprague-Dawley rats that progressed to compulsive stimulant use [86], highlighting a critical role for trait compulsivity, and perhaps prefrontal cortex and nucleus accumbens circuit functions, in the development of cocaine addiction.

### 4.4 Limitations

This study has some limitations. First, only male rats were included in this feasibility study. Given our limited resources and the logistical difficulties in running a large sample size in a longitudinal deep phenotyping study, as a first step, we limited the evaluation to males in order to facilitate comparisons with previous studies which by and large, employed males-only. We fully acknowledge the importance of running both sexes and will employ both females and males in future replication studies and in an F2 cross. Second, the h^2^ heritability estimates are specific for the phenotypes we measured in parental SHR/NCrl and SHR/NHsd substrains.

More precise estimates will be obtained in the F2 cross, including SNP h^2^ estimates. Lastly, compulsive cocaine use, defined as continued cocaine seeking despite aversive consequences, needs to be studied. This definitive clinical and preclinical measure of cocaine addiction will require much larger group sizes than what was used herein because resistance to punishment only occurs in a subset of rats (∼25%) that receives extended self-administration training [41].

This specific phenotype can more easily be assessed in an F2 cross, as large sample sizes are required for identifying the multiple genes of small effect size that contribute to complex traits [11].

## 5. Conclusions

These results provide compelling support for using an RCC approach in SHR substrains to uncover candidate genes and variants that are of relevance to cocaine use disorders. The robust substrain differences in multiple premorbid addiction vulnerability traits and cocaine self-administration behaviors support the heritability of these traits and increase the likelihood that a forward genetic mapping approach will be successful. The advantage of an RCC is that this type of cross segregates orders of magnitude fewer genetic variants, making an RCC a simple and powerful solution for rapid, high-confidence gene discovery for complex traits [11].

## Author Contributions

**Kathleen Kantak:** Funding Acquisition, Conceptualization, Methodology, Formal Analysis, Visualization, Writing. **Carissa Stots and Elon Mathieson:** Investigation and Data Curation, **Camron Bryant:** Funding Acquisition, Conceptualization, Methodology, Formal Analysis, Visualization, Writing.

